# Imaging increased metabolism in the spinal cord in mice after middle cerebral artery occlusion

**DOI:** 10.1101/2022.08.11.503550

**Authors:** Ruiqing Ni, Nadja Straumann, Serana Fazio, Xose Luis Dean-Ben, Georgios Louloudis, Claudia Keller, Daniel Razansky, Simon Ametamey, Linjing Mu, César Nombela-Arrieta, Jan Klohs

**Affiliations:** Institute for Regenerative Medicine, University of Zurich, Zurich, Switzerland; Institute for Biomedical Engineering, Department of Information Technology and Electrical Engineering, University of Zurich & ETH Zurich, Zurich, Switzerland; Zentrum für Neurowissenschaften Zurich, Zurich, Switzerland; Department of Medical Oncology and Hematology, University and University Hospital Zurich, Zurich, Switzerland; Center for Radiopharmaceutical Sciences ETH, PSI and USZ, Institute of Pharmaceutical Sciences, Department of Chemistry and Applied Biosciences, ETH Zurich, Zurich, Switzerland

**Keywords:** fluorodeoxyglucose (FDG), ischemic stroke, metabolism, optoacoustic tomography, positron emission tomography, spleen, spinal cord, Ischemia, metabolic activity, MSOT, oxygenation, PET

## Abstract

Emerging evidence indicates crosstalk between the brain and the hematopoietic system following cerebral ischemia. Here, we investigated metabolism and oxygenation in the spleen and spinal cord in a transient middle cerebral artery occlusion (tMCAO) mouse model that is widely used in focal cerebral ischemia research. Naïve, sham and tMCAO mice underwent positron emission tomography (PET) using [^18^F]fluorodeoxyglucose (FDG) for assessing glucose metabolism and multispectral optoacoustic tomography (MSOT) assisted with quantitative model-based reconstruction and unmixing algorithms for accurate mapping of oxygenation patterns in the peripheral tissues at 24 h after reperfusion. We found increased levels of [^18^F]FDG uptake and reduced MSOT oxygen saturation, indicating hypoxia in the thoracic spinal cord of tMCAO mice compared with sham-operated mice but not in the spleen. A positive correlation was observed between splenic and ipsilateral striatal [^18^F]FDG uptake. Reduced spleen size was observed in tMCAO mice compared with sham-operated mice *ex vivo*. tMCAO led to a significant increase in the numbers of mature T cells (CD4 and CD8) in femoral bone marrow tissues, concomitant with a stark reduction in these cell subsets in the spleen and their decrease in peripheral blood. The numbers of mature granulocytes (determined as CD11b^+^Gr1^hi^ cells) decreased in bone marrow tissues and blood but increased in the spleen. The combination of quantitative PET and MSOT thus enabled the observation of hypoxia and increased metabolic activity in the spinal cord of tMCAO mice at 24 h after occlusion compared to sham-operated mice.

## 1. Introduction

Brain injury due to cerebral ischemia results in early activation of the immune system that is at a later stage followed by immunodepression [1, 2]. In experimental cerebral ischemia, the inflammatory response is crucially involved in secondarily enhancing lesion growth but is also indispensable for tissue repair and regeneration. Immunodepression after cerebral ischemia increases the susceptibility to infections [3], leading to impaired recovery and death. Thus, interactions between the brain and the immune system are very relevant to the overall outcome after cerebral ischemia [4], and understanding these processes is key before immunomodulatory therapy can be applied to stroke patients.

Levels of circulating neutrophils and monocytes increase in the circulation [5, 6]. These myeloid cells are recruited to the ischemic brain [7, 8]. Monocytes are mobilized from the spleen, which is the largest lymphatic organ in the body, upon injury [9]. Altered splenic function, reductions in spleen weight and a decrease in monocytes/macrophages in the spleen have been demonstrated in both permanent and transient middle cerebral artery occlusion (pMCAO and tMCAO) models [10–14]. Moreover, it was shown that tMCAO activates bone marrow (BM) hematopoietic stem cells (HSCs) and downstream hematopoietic progenitors, leading to an increased output of inflammatory monocytes and neutrophils [15]. Activation of the hematopoietic organs is an active area of research, and studies have shown that progenitor and immune cell function are controlled by tissue oxygen tension and glucose metabolism [16, 17]. However, few studies have noninvasively investigated metabolic processes in hematopoietic organs in experimental models of cerebral ischemia and stroke patients.

Several studies have applied [^18^F]fluorodeoxyglucose (FDG) PET to assess glycolysis of hematopoietic organs in patients with a variety of disease conditions, including infection [18], cancer [19], vasculitis [20], and cardiovascular diseases [21, 22]. Studies have found marked elevations in [^18^F]FDG uptake in the spleen and spinal cord, which was associated with systemic inflammatory markers and future cardiovascular events. Studies have demonstrated decreased [^18^F]FDG uptake in lumbar vertebrae (L4, L5), and no difference in the spleen of ischemic stroke patients compared to controls has been reported, but PET was performed at the chronic stage of the disease with a larger variability [23]. [^18^F]FDG PET of hematopoietic organs in MCAO models, where the assessment of earlier time points after cerebral ischemia with lower temporal variability is easier to attain, has thus far not been performed.

Another regulator of immune and stem cell function in hematopoietic organs is local tissue oxygen tension [17]. Optoacoustic imaging allows for the non-invasive assessment of blood oxygenation relatively deep in tissue (mm to cm range) [24–28]. It has been previously employed to detect functional and structural alterations in the brain but not yet in the periphery of tMCAO animal models [29–33]. In this study, we aimed to image metabolic alterations in the hematopoietic organs in response to tMCAO with [^18^F]FDG PET and multi-spectral optoacoustic tomography (MSOT) and evaluate immune and progenitor cell responses using flow cytometry.

## 2. Materials and methods

### 2.1 Animal model

Twenty-four male C57BL/6J mice (Janvier, France) weighing 20-25 g and 8-10 weeks of age were used. Mice were randomly allocated to naïve (n = 3), sham (n = 10) or tMCAO (n = 11) groups. Animals were housed in ventilated cages inside a temperature-controlled room under a 12 h dark/light cycle. Pelleted food (3437PXL15, CARGILL) and water were provided *ad libitum*. Paper tissue and red mouse house® (Tecniplast, Milan, Italy) shelters were placed in cages for environmental enrichment. All experiments were performed in accordance with the Swiss Federal Act on Animal Protection and were approved by the Cantonal Veterinary Office Zurich (permit number: ZH080/18). We confirm compliance with the NC3Rs ARRIVE 2.0 guidelines on reporting of in vivo experiments.

### 2.2 tMCAO

Surgeries for tMCAO and sham operation were performed as described previously [34, 35]. Anaesthesia was initiated by using 3 % isoflurane (Abbott, Switzerland) in a 1:4 oxygen/air mixture and maintained at 2 %. In addition, buprenorphine was administered as subcutaneous (s.c.) injection (0.1 mg/kg; 1 ml Temgesic + 5 ml NaCl, inject 2 μl Temgesic/g of body weight). The temperature was kept constant at 36.5 ± 0.5 °C with a feedback-controlled warming pad system. All surgical procedures were performed in less than 15 min. After surgery, a second dose of buprenorphine was administered as s.c. injection (0.1 mg/kg; 1 ml Temgesic + 5 ml NaCl, inject 2 μl Temgesic/g of body weight) 4 h after reperfusion and supplied thereafter by drinking water (0.1 mg/kg body weight) until the end of the study. Animals received softened chow in a weighing boat on the cage floor to encourage eating. MCAO animals were excluded from the study if they met one of the following criteria: Bederson score of 0, no reflow after filament removal, and premature death.

### 2.3 [^18^F]FDG-μPET/CT

Four sham and 4 tMCAO mice underwent microPET imaging at 24 h after reperfusion. Scans were performed with a calibrated SuperArgus μPET/CT scanner (Sedecal, Spain) with an axial field-of-view of 4.8 cm and a spatial resolution of 1.6-1.7 mm (full width at half maximum). tMCAO and sham-operated C57BL/6J mice were anesthetized with isoflurane (2.5 %) in oxygen/air (1:1) during tracer injection and the whole scan time period. The formulated radioligand solution ([^18^F]FDG: 9.9-11 MBq) was administered via tail vein injection. Mice were dynamically scanned until 60 min after injection. Body temperature was monitored by a rectal probe and kept at 37 °C by a heated air stream (37 °C). The anesthesia depth was measured by respiratory frequency (SA Instruments, Inc., USA). μPET acquisitions were combined with CT for anatomical orientation and attenuation correction. The obtained data were reconstructed in user-defined time frames with a voxel size of 0.3875×0.3875×0.775 mm^3^ as previously described [36].

### 2.4 PET data analysis

Images were processed using PMOD 4.2 software (PMOD Technologies Ltd., Zurich, Switzerland). Spleen, cervical and thoracic spinal cord volumes of interest (VOIs) that were defined based on the CT contrast. The time-activity curves were deduced from specific VOIs. Radioactivity is presented as the standardized uptake value (SUV) (decay-corrected radioactivity per cm^3^ divided by the injected dose per gram body weight). The SUV signal was averaged from 18-53.5 minutes. Vertebral VOIs were drawn manually similar to a previous report (**SFig. 1**) [37].

### 2.5 MSOT

Three naïve, six sham and four tMCAO mice underwent MOST imaging at 24 h after reperfusion. An MSOT inVision 128 imaging system (iThera Medical, Germany) was used as described previously [38–40]. Briefly, a tunable (680-980 nm) optical parametric oscillator pumped by a Nd:YAG laser provides <10 ns excitation pulses at a frame rate of 10 Hz with a wavelength tuning speed of less than 10 ms and a peak pulse energy of 100 mJ at 730 nm. Ten arms, each containing an optical fiber bundle, provide even illumination of a ring-shaped light strip with a width of approx. 8 mm. For ultrasound detection, an array of 128 cylindrically focused ultrasound transducers with a center frequency of 5 MHz (60 % bandwidth), organized in a concave array of 270 degree angular coverage and a curvature radius of 4 cm, were used.

For *in vivo* MSOT measurement, animals were anesthetized with 4 % isoflurane and maintained at 1.5 % isoflurane in a 1:4 oxygen/air mixture supplied via a nose cone. Mice were depilated around the body parts of interest (approximately below the 5^th^ rib) and placed in a mouse holder in the supine position with a cuff around the legs. Preheated ultrasound gel (Diagramma, Switzerland) was applied to the mouse body for ultrasonic coupling, and the animals were wrapped in a polyethylene membrane. The mouse holder was placed in an imaging chamber filled with water with temperature controlled at 36.5 °C. Images were acquired at 5 wavelengths (715, 730, 760, 800, and 850 nm) for 135-140 consecutive slices with a step size of 0.3 mm and 10 averages. The total acquisition time was approximately 10 minutes.

### 2.6 MSOT image reconstruction and spectral unmixing

To avoid negative-value artifacts present in the images reconstructed with standard back projection algorithms [26]. MSOT images were processed with a dedicated non-negative constrained model-based (NNMB) reconstruction-unmixing framework [41], which is also known to facilitate quantification of oxygenation values in the images [43]. Specifically, images corresponding to different excitation wavelengths were first reconstructed with an NNMB algorithm. Subsequently, linear unmixing was also performed with an NNMB algorithm considering the theoretical absorption spectra of deoxygenated (Hb) and oxygenated (HbO_2_) hemoglobin. For comparison, the images were also processed with the data analysis system software (iTheraMedical, Germany). This consists of a standard model-based reconstruction algorithm followed by linear spectral unmixing but not including non-negative constraints. Unmixed images represent the biodistribution of Hb and HbO_2_. The oxygen saturation (sO_2_) of the tissue (the capillaries/microvasculature composing the tissue) was then calculated as

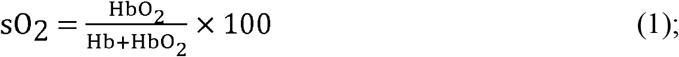

Regions of interest (ROIs) for the spleen, thoracic spinal cord and femur were drawn manually on the sO_2_ map to the structural reference (MSOT image corresponding to 850 nm excitation) using ImageJ (NIH, USA). Line profile analysis was performed to compare the signal intensity of different reconstruction methods. Regions-of-interest (ROIs) were drawn manually over the Z-stack for the spleen and femur (on the left and right hindlimb). At least three independent ROI delineations over 30-40 horizontal slices were performed. Volumes were calculated considering the known slice gap during image acquisition (0.3 mm). The results represent the averages of all analyses.

### 2.7 FACS analysis of BM, spleen and blood

Murine femurs were harvested through careful dislocation at the joints between the hipbones and the knee and cleaned thoroughly of surrounding connective tissue and muscle using paper tissues. Mouse calvarial bones were exposed by making an incision at the base of the neck and removing the skin flap of the skull. Using microdissection scissors, the skull bone was then cut from the neck’s base towards the nose bridge and carefully peeled from the underlying brain tissue. Murine spleens were extracted by abdominal incision. All organs were preserved in FACS buffer on ice before further processing. In turn, blood was collected immediately after euthanasia via intracardiac puncture using a 5 ml syringe and a 25 G needle. The samples were immediately transferred into a potassium ethylenediamine tetra-acetic acid (K+ EDTA) blood collection tube and shaken vigorously for 15 seconds to avoid coagulation.

For the generation of single-cell suspensions, bones were crushed in 5 mL of FACS buffer (phosphate buffered saline with 2 % fetal bovine serum and 2 mM EDTA), and the released cellular content was disaggregated via resuspension using a pipette. The cell suspensions were then filtered through a 70 μm cell strainer into a 50 mL collection tube. Bones were crushed as described three times to maximize marrow extraction and until bone fragments appeared completely pale. Spleens were crushed in FACS buffer directly into a 70 μm cell strainer using a syringe plunger under repeated washing. For the BM and spleen, red blood cells (RBCs) were removed by lysis in 2 mL of RBC lysis buffer (BioLegend) for 3 minutes. In the case of blood, 100 μl samples were lysed on 3 ml of RBCs for 3 min, and the reactions were inactivated through the addition of excess FACS buffer. Cells were then centrifuged, washed, and filtered once more before immunostaining for FACS analysis. Cells were then resuspended in fixed volumes, and the samples were blocked using Fc Receptor Blocking Solution (TruStain FcX^tm^) at 4 °C. Cocktails of fluorescently labelled antibodies were added (**Supplementary Table 1**), and the cells were incubated for 30 minutes at 4 °C. Cells were then washed twice in FACS buffer, resuspended in FACS buffer containing 4’,6-diamidino-2-phenylindole (DAPI; 0.5 ug/mL), and analysed on a LSR II Fortessa (BD Biosciences). Data analysis was performed using FlowJo software, version 10.8.

### 2.8 2,3,5-triphenyltetrazolium chloride (TTC) staining and spleen size and mass measurement

Staining with 2,3,5-triphenyltetrazolium chloride (TTC) was used to assess ischemic lesion severity as previously described [36]. Mice were euthanized, their brains were removed, and 1 mm thick brain slices were obtained with a brain matrix (World Precision Instruments). Slices were incubated in a 2.5 % TTC solution (Sigma–Aldrich, Switzerland) in 1×phosphate-buffered saline (PBS, pH 7.4) at 37 °C for 3 min. Photographs of the brain sections were taken. The spleens were dissected and measured using a ruler and weighed.

### 2.9 Statistics

For data analysis, GraphPad Prism (GraphPad Prism 9.2.0, USA) and RStudio (R Core Team, Austria) [42] were used. An unpaired two-tailed t test was used for the comparison of regional [^18^F]FDG SUV between the sham and tMCAO groups. One-way ANOVA with Sidak post hoc analysis was used for MSOT-derived sO_2_, splenic volume and mass comparisons between the three groups. Two-way ANOVA was used to compare the time course between groups. Spearman rank correlation analysis was used to analyse the link between the [^18^F]FDG SUV and sO_2_ in the brain, spinal cord and spleen. Data are presented as the mean ± standard deviation (SD). Significance was set at p < 0.05.

## 3. Results

### 3.1 Focal cerebral ischemia leads to increased thoracic spinal cord metabolism

To investigate metabolic activity after the hematopoietic tissues after focal cerebral ischemia, tMCAO and sham-operated mice underwent [^18^F]FDG-μPET (**Fig. 3a, b**). The time activity curves of [^18^F]FDG uptake in the spleen and spinal cord of the sham and tMCAO groups are shown in **Figs. 3c, h**, and **SFig. 1**. We found an approximately 20 % increase in [^18^F]FDG uptake (average 18-53.5 minutes) in the spinal cord (C1-T13) of tMCAO mice (0.16 ± 0.02, n = 4) compared to sham-operated mice (0.13 ± 0.01, n = 4, p = 0.032) (**Fig. 1d**). Next, we further quantified [^18^F]FDG uptake in cervical (C1-7) and thoracic (T1-13) spinal cord segments (**Figs. 1e, f**). [^18^F]FDG uptake (averaged 18-53.5 minutes) was not different in the cervical part of the spinal cord of tMCAO mice (0.18 ± 0.03, n = 4) compared to sham-operated mice (0.14 ± 0.01, p = 0.0516) (**Fig. 1e**). In contrast, higher [^18^F]FDG uptake (averaged 18-53.5 minutes) was observed in the thoracic part (T1-13) of tMCAO mice (0.151 ± 0.02, n = 4) than in sham-operated mice (0.124 ± 0.004, p = 0.0384) (**Fig. 1f**). In addition, [^18^F]FDG uptake in the cervical part was higher than that in the thoracic part (0.16 ± 0.03 vs 0.14 ± 0.02, p = 0.0004) (**Fig. 1g**). There were no differences in splenic [^18^F]FDG SUV uptake (averaged 18-53.5 minutes) in tMCAO mice (1.23 ± 0.01, n = 4) and sham-operated mice (1.22 ± 0.02, n = 4) at 24 h after occlusion (**Fig. 1i**). To examine whether there is a potential link between the brain and peripheral [^18^F]FDG uptake, Spearman rank analysis was performed between different readouts. There was a positive correlation between the ipsilateral striatal and splenic [^18^F]FDG SUV (averaged 18-53.5 minutes, p = 0.0011, r = 0.9524 Spearman rank analysis). No correlation was observed between ipsilateral striatal and spinal cord [^18^F]FDG SUV (**Fig. 1j, k**).

**Fig. 1.**
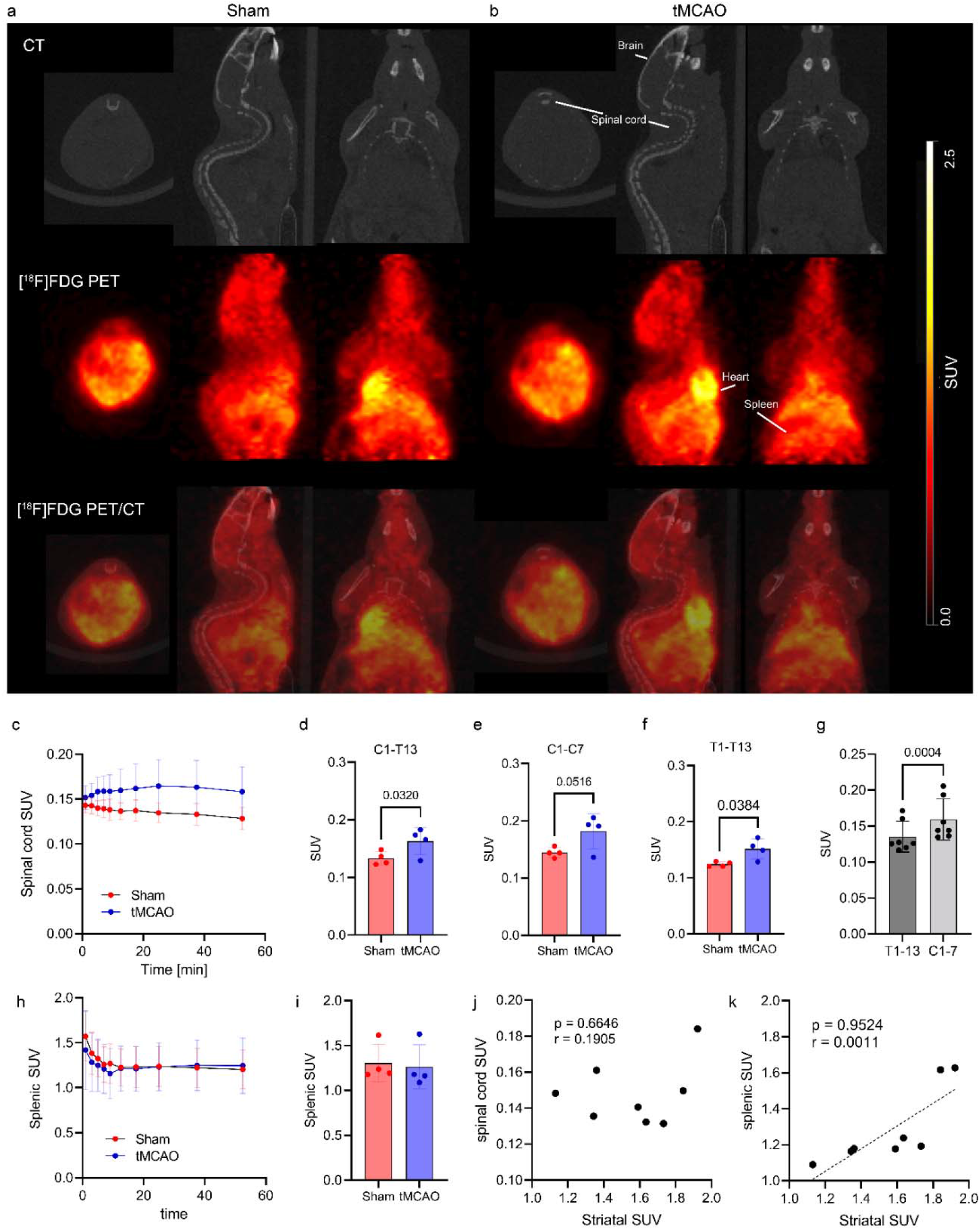
*In vivo* [^18^F]FDG PET of metabolism in the spleen of sham and tMCAO mice at 24 h after occlusion. (**a, b**) Representative coronal, sagittal and horizontal views of [^18^F]FDG uptake (Bq/ml) in sham-operated and tMCAO mice (n = 4 per group, averaging 18-53.5 min); CT, computed tomography. (**c, h**) Time activity curve of [^18^F]FDG in the spinal cord and spleen of sham-operated and tMCAO mice. The volumes of interest for the spleen and spinal cord were manually delineated (white line for spleen). Difference in the spinal cord TAC between sham and tMCAO (mixed effect model treatment, p = 0.0399, treatment x time p = 0.0073), (**d-f**) The [^18^F]FDG SUV signal was higher in the spinal cord (C1-T13, and C1-7, T1-13) of tMCAO mice compared to sham-operated mice. (**g**) The SUV is higher in C1-7 than in T1-13. (**i**) No difference in [^18^F]FDG SUV (n = 4 per group, averaging 18-53.5 min) in the spleen FDG in sham-operated and tMCAO mice was detected. (**j-k**) No correlation between striatal and spinal cord [^18^F]FDG SUV. (**i**) Positive correlation between striatal and splenic [^18^F]FDG SUV (Spearman rank analysis). Data are presented as the mean ± standard deviation (SD).

### 3.2 NNMB reconstruction and unmixing

We used the MSOT system (**Fig. 2a**) for non-invasive visualization of the mouse body. The MSOT imaging pipeline consisted of reconstructing 3D images acquired at multiple excitation wavelengths (**Fig. 2**) and spectral unmixing for different tissue chromophores, followed by manual VOI analysis. The reconstruction-unmixing processing steps are described in the methods section. The enhanced quantitative performance achieved with the NNMB method has been previously demonstrated in phantom studies [41] and in brain imaging data (orthotopic glioblastoma model) [43]. We compared the signal intensities of the images obtained with the custom-made NNMB reconstruction-unmixing framework with those of the images obtained with the standard MB approach integrated in the MSOT software package (**Figs. 2b-h**). The application of NNMB processing (reconstruction and unmixing) facilitated enhancing image features corresponding, e.g., to the spinal cord and the spleen as well as to other organs at deeper locations in the structural images and, to a greater extent, in the sO_2_ images (indicated by the line profile analysis) (**Figs. 2i, j**).

**Fig. 2.**
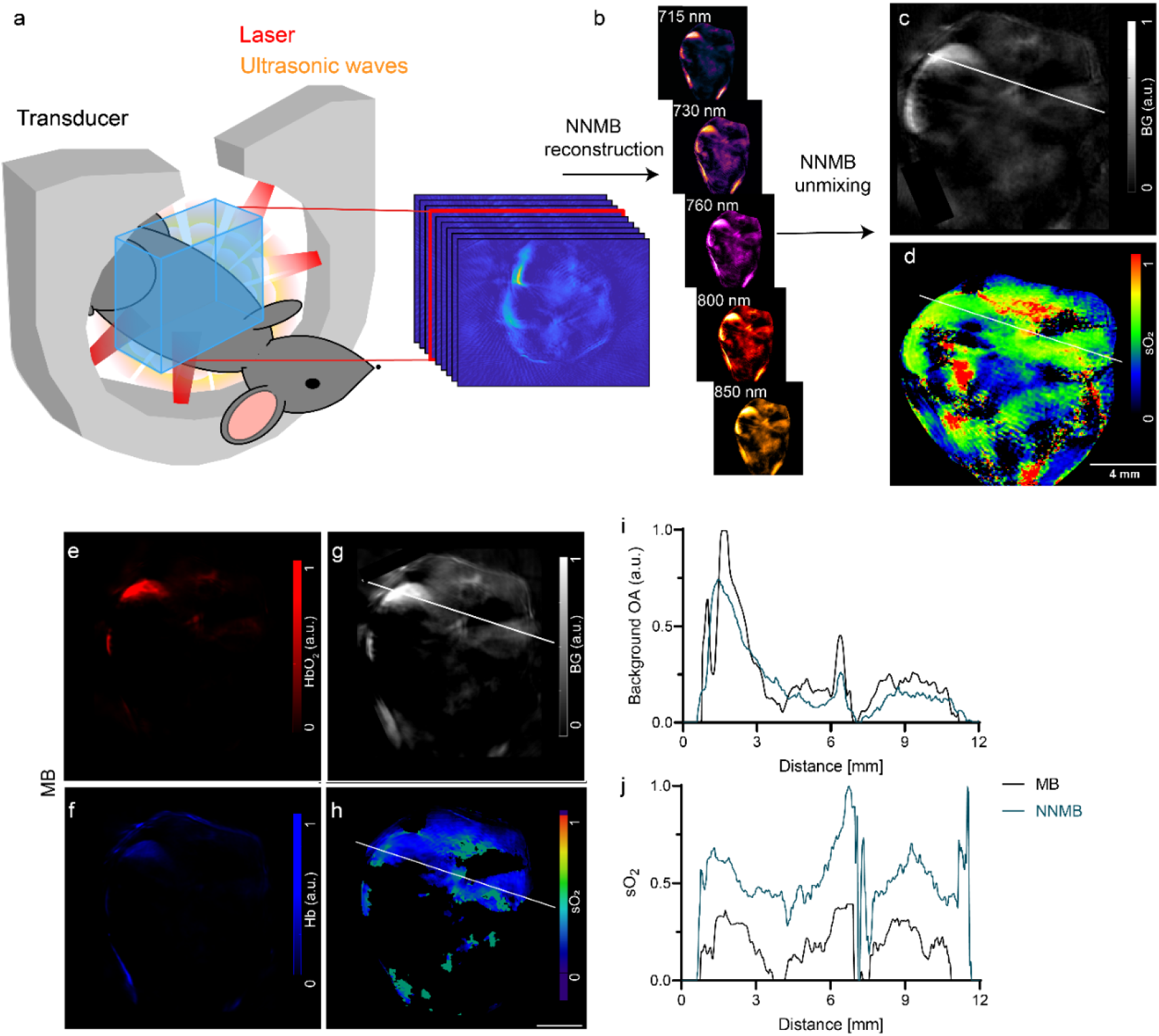
*In vivo* MSOT of the mouse body and data processing. (**a**) Setup of the MSOT system for imaging across the mouse body; (**b**) non-negative model-based (NNMB) reconstructions of the *in vivo* vMSOT data for five distinct excitation wavelengths (715-850 nm). (**c, d**) Horizontal views of a background anatomy and oxygen saturation rate (sO_2_) slice resulting from NNMB unmixing. Scale bar = 4 mm. (**e-h**) Same set of image data after standard model-based (MB) reconstruction and unmixing, HbO_2_ (red), Hb (blue), background anatomy, and sO_2_ slice. (**i-j)** Profile analysis of c, d and g, h showed differences in the background OA intensity and in sO_2_ values processed by the MB and NNMB methods.

**Fig. 3.**
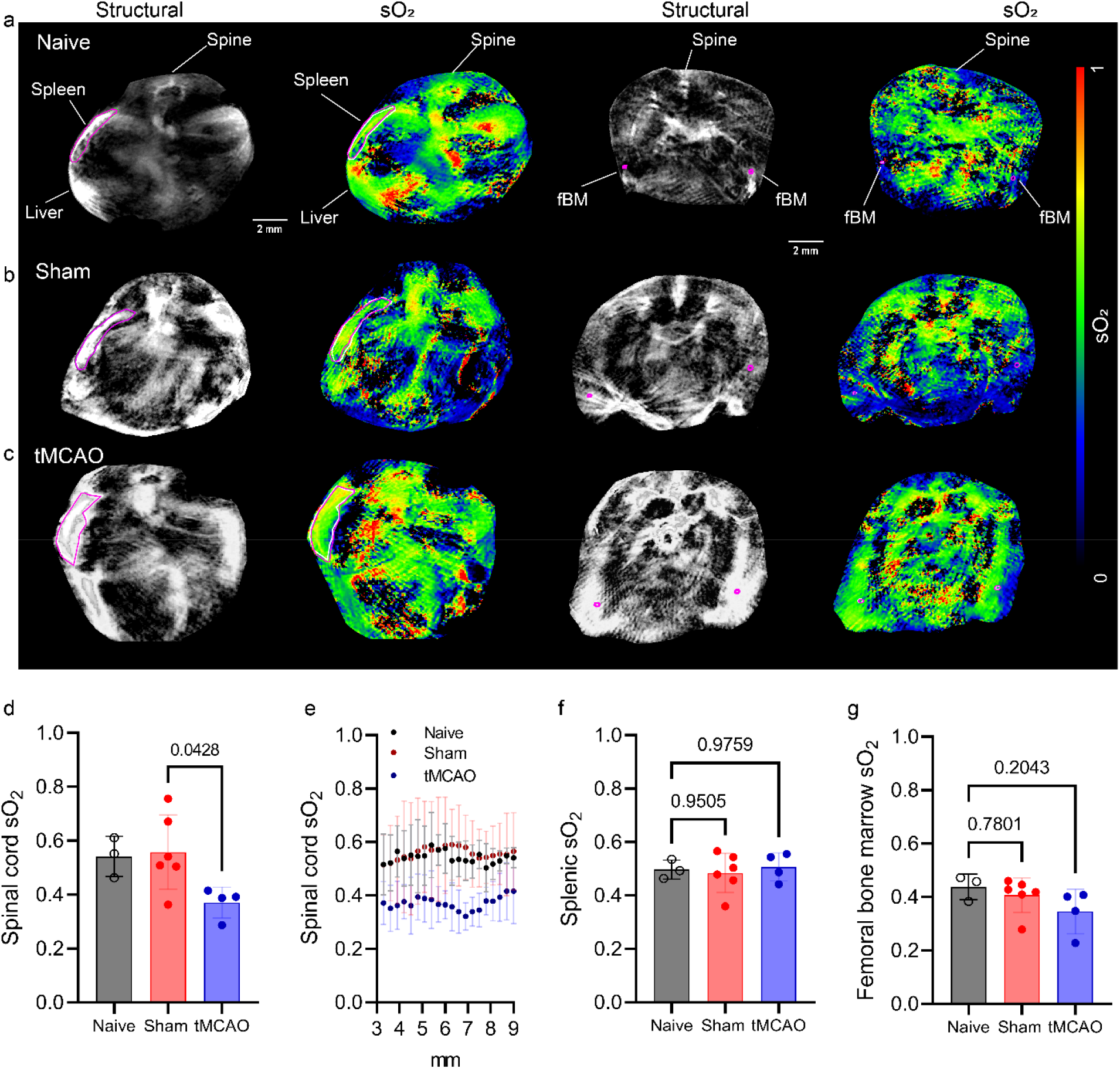
*In vivo* MSOT of hemodynamics in the spinal cord, spleen and femur of naïve, sham and tMCAO mice at 24 h after occlusion. (**a-c**) Oxygen saturation rate (sO_2_) map in the spinal cord, spleen and femoral bone marrow (fBM) of naïve, sham-operated and tMCAO mice. (**d, f, g**) Reduced spinal cord oxygen saturation rate (sO_2_) and comparable in the spleen and femoral bone marrow (fBM) of naïve (n = 3), sham-operated (n = 6) and tMCAO mice (n = 4). (**e**) Profile of sO_2_ across different positions of the spinal cord. Data are presented as the mean ± standard deviation (SD).

### 3.3 Focal cerebral ischemia leads to reduced thoracic spinal cord oxygenation

To reveal oxygenation in the thoracic spinal cord and spleen after tMCAO *in vivo*, mice were assessed with MSOT at 24 h after occlusion. The spleen, cervical and thoracic spinal cord and femur in the representative structural map resulting from NNMB processing had clear boundaries and were used for ROI delineation and later transferred to sO_2_ images for data analysis (**Fig. 3a-c**). In the thoracic spinal cord, the sO_2_ of the tMCAO group (0.37 ± 0.06, n = 4) was lower than that of the sham-operated group (0.56 ± 0.14, n = 6, p = 0.0428) (**Fig. 3d**). Variation in the sO_2_ across the thoracic spinal cord was observed. Comparable sO_2_ was detected in the spleen of naïve (0.5 ± 0.04, n = 3), sham-operated (0.5 ± 0.07, n = 6), and tMCAO mice (0.5 ± 0.05, n = 4) (**Fig. 3f**) and in the femur of naïve (0.4 ± 0.05, n = 3), sham-operated (0.5 ± 0.07, n = 6), and tMCAO mice (0.3 ± 0.08, n = 4) (**Fig. 3g**).

### 3.4 Alterations in hematopoietic and lymphoid organs after tMCAO

Flow cytometry analysis revealed major changes in hematopoietic and lymphoid organs, as well as circulating cells, after tMCAO and sham surgeries. In general, we observed that tMCAO led to a significant increase in the numbers of mature T cells (CD4 and CD8) in femoral BM tissues (**Figs. 4e, f**), concomitant with a stark reduction in these cell subsets in the spleen (**Figs. 5e, f**) and their decrease in peripheral blood (**Figs. 6d, e**). In turn, the numbers of mature granulocytes (determined as CD11b^+^Gr1^hi^ cells) decreased in BM tissues (**Fig. 4b**) and blood (**Fig. 6a**) but increased in the spleen (**Fig. 5b**). Of note, both trends were also observed to a large extent in sham-operated mice. Notably, as previously reported, tMCAO induced subtle but significant increases in the frequencies and absolute numbers of hematopoietic progenitor cells (Lin^-^c-kit^+^Sca-1^+^, LSK) (**Figs. 4g, n**) and HSCs (LSKCD48^-^ CD150^+^) (**Figs. 4h, o**). Again, alterations in these early primitive hematopoietic progenitors were also found, albeit at lower magnitudes, in sham controls (**Figs. 4c, j**). This suggests that the profound redistribution of mature cell types from peripheral circulation and spleen to BM tissues as well as the reactive expansions of early progenitor populations and HSCs are largely caused by surgery-associated stress and accentuated in mice subjected to tMCAO.

**Fig. 4.**
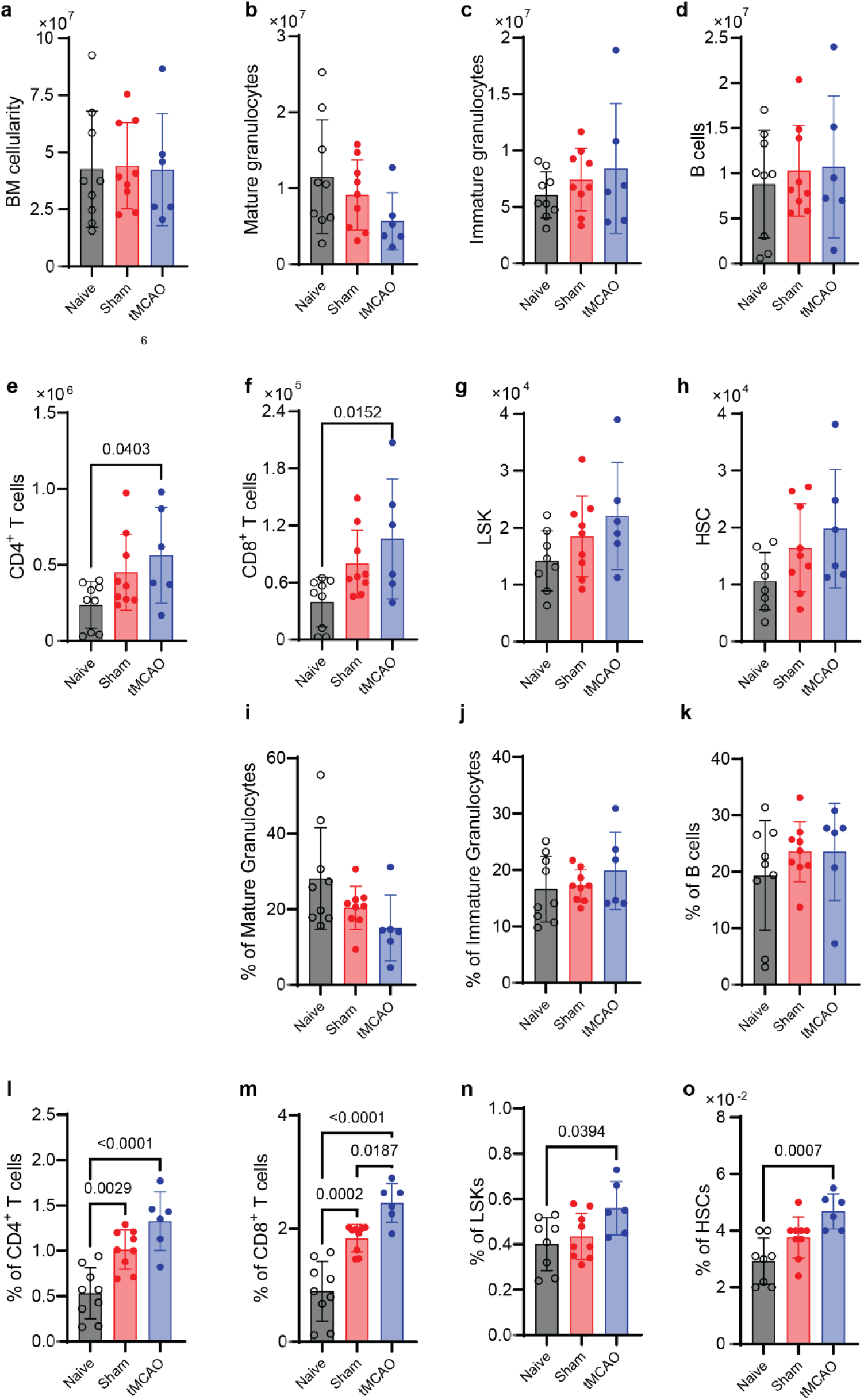
tMCAO increased the numbers of mature T cells in femur bone marrow. Flow cytometric analysis of the absolute number **(a-h)** and percentage **(i-o)** of mature granulocytes, immature granulocytes, B cells, CD4^+^ T cells, CD8^+^ T cells, LSCs and HSCs in the femur bone marrow of naïve, sham and tMCAO mice (n = 6-9 per group).

**Fig. 5.**
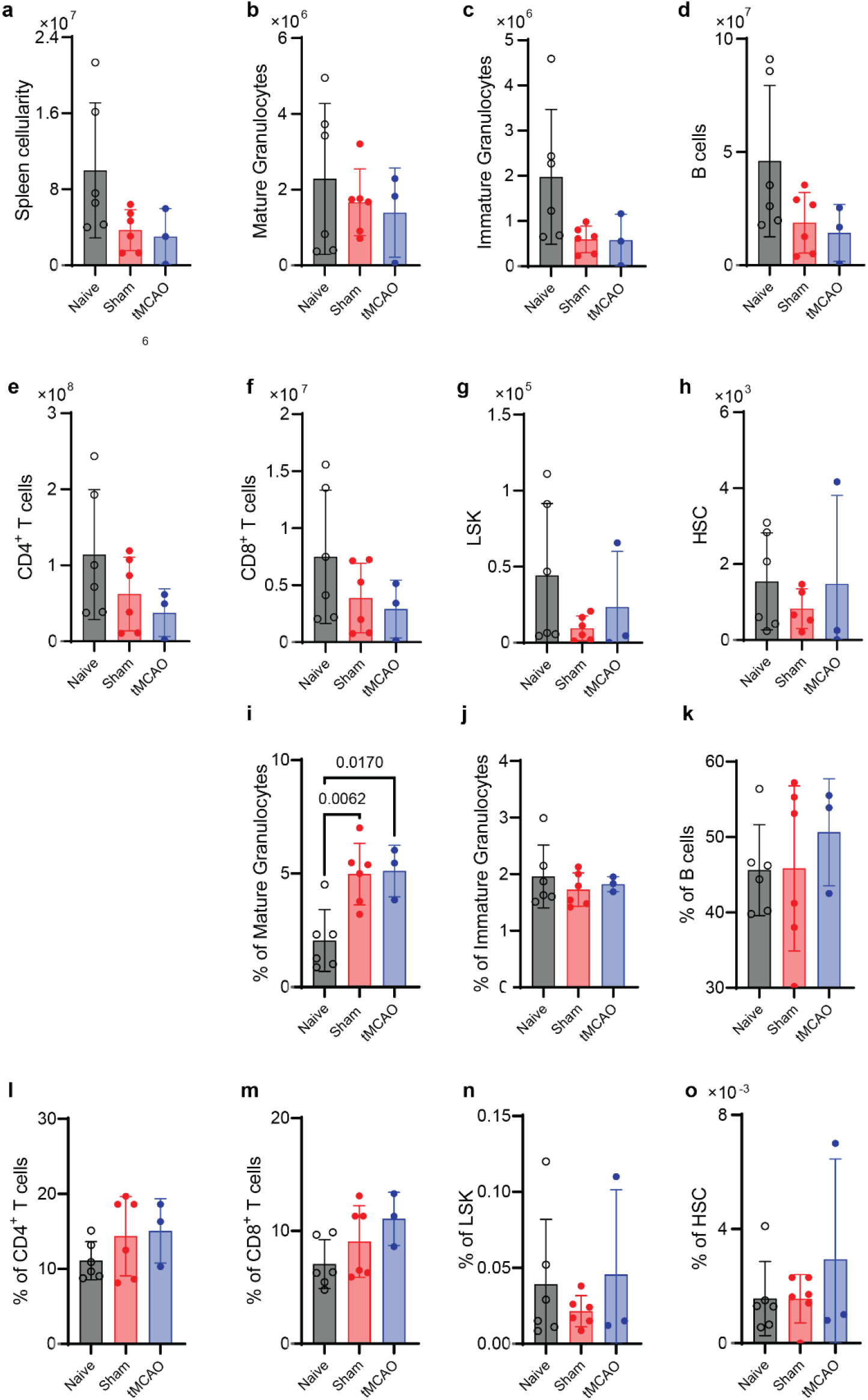
tMCAO increased mature granulocytes in the spleen. Flow cytometric analysis of the absolute number **(a-h)** and percentage **(i-o)** of mature granulocytes, immature granulocytes, B cells, CD4^+^ T cells, CD8^+^ T cells, LSCs and HSCs in the spleen of naïve, sham and tMCAO mice (n = 6-9 per group).

**Fig. 6.**
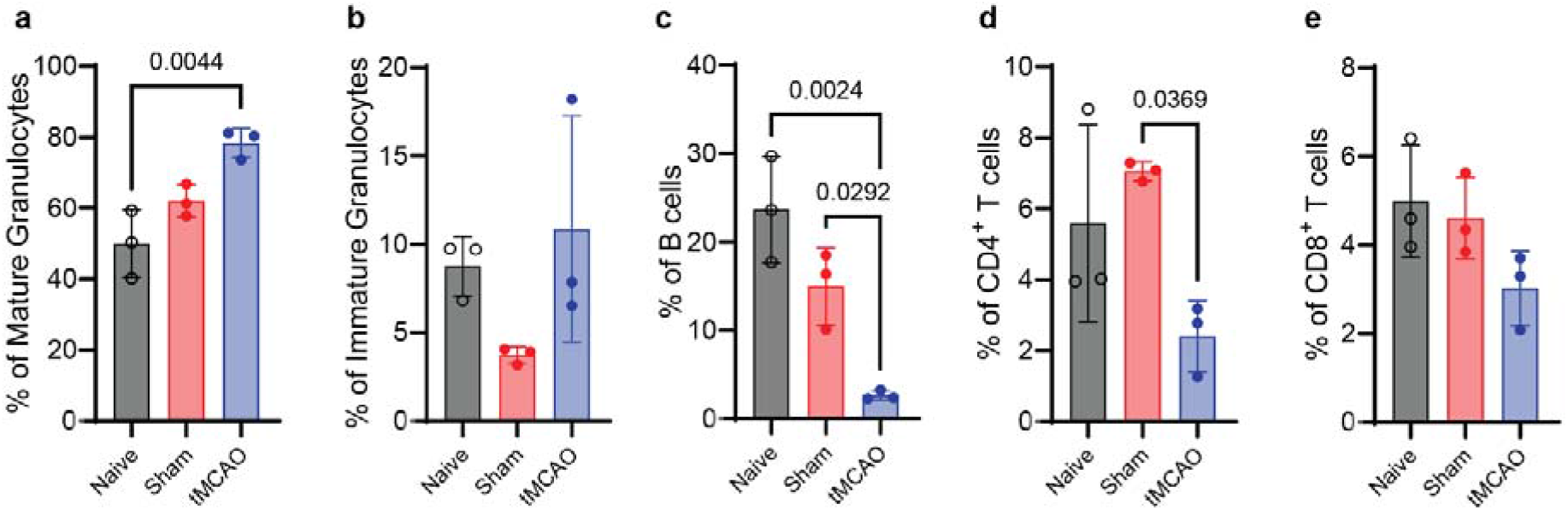
tMCAO reduced B cells and T cells in circulating blood. **(a-e)** Flow cytometric analysis of the percentage of mature granulocytes, immature granulocytes, B cells, CD4+ T cells and CD4+ T cells in the circulating blood of naïve, sham and tMCAO mice (n = 3 per group).

### 3.5 Histology and TTC staining validation and spleen pathology

Significantly reduced weight in both sham-operated (22.1 ± 2.1 g, p = 0.0425) and tMCAO mice (21.6 ± 2.0 g, p = 0.0054) was detected compared to naïve mice (25.6 ± 1.3 mg, p = 0.0029). Ischemic lesions in the tMCAO mice were validated by the presence of nonviable tissue white areas shown by TTC staining (approximately 42 %, **Figs. 7a, d**). The spleens of the sham-operated and tMCAO mice were dissected, and the splenic size and mass were measured after *in vivo* imaging (**Figs. 7b, c**). Compared to naïve mice (88.7 ± 31.7 mg, n = 6), reduced splenic mass at approximately 25 h after reperfusion (after *in vivo* imaging) was detected in both the sham and tMCAO mice (**Fig. 7e**). No difference in the *ex vivo* splenic size was detected between sham and tMCAO mice.

**Fig. 7.**
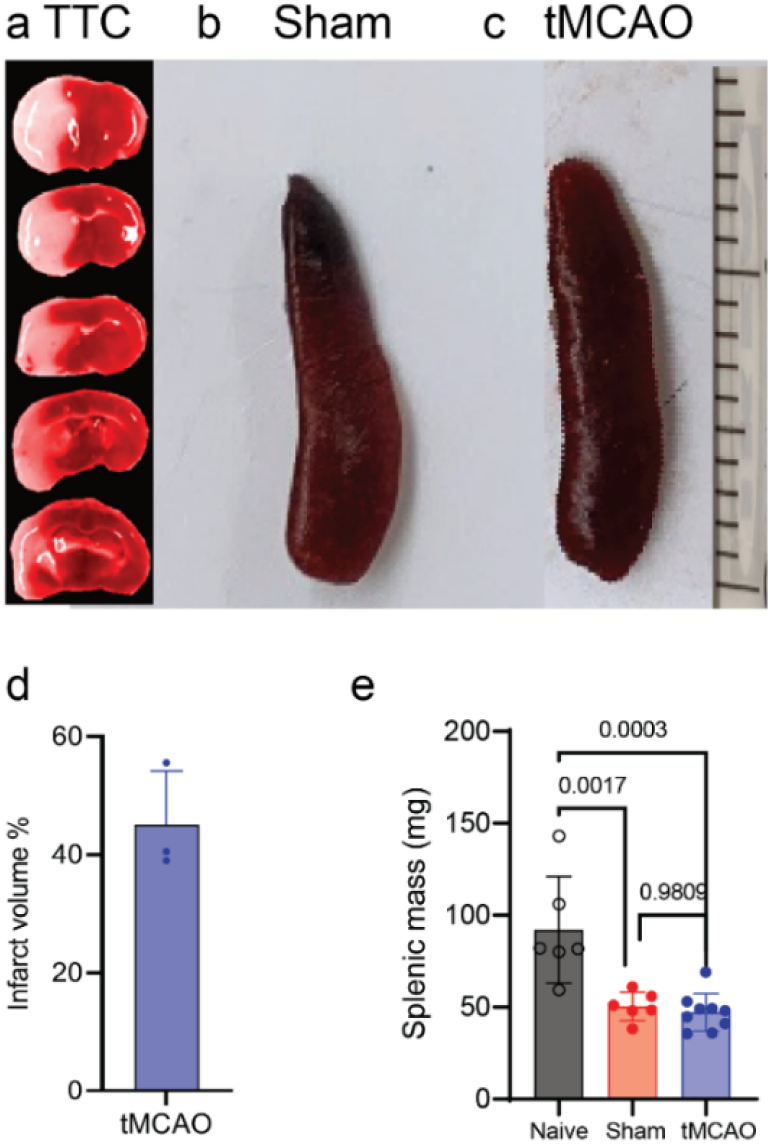
Assessment of the spleen volume Histology of the brain and spleen of sham and tMCAO mice at 24 h after occlusion. (**a**) TTC of tMCAO mouse brain slices indicating lesions in the ipsilateral striatum and cortex. (**b, c**) Representative images of the dissected spleen from sham-operated and tMCAO mice. (**d**) Percentage of infarct volume in the brains of tMCAO mice. (**e**) Reduced mass of dissected spleen among naïve (n = 6), sham-operated (n = 6) and tMCAO mice (n = 8) *ex vivo*. Data are presented as the mean ± standard deviation (SD).

## 4. Discussion

Emerging evidence indicates brain-periphery crosstalk and activation of the hematopoietic system after ischemic stroke via the sympathetic nervous system [15]. tMCAO induces rapid activation of the peripheral immune system in the spleen and spinal cord as well as blood–spinal cord barrier alterations in rodent and primate models [11, 44–48]. Here, we demonstrated non-invasive imaging of increased metabolism and reduced oxygen saturation in the spinal cord but not in the spleen of tMCAO mice at 24 h after 1 h of occlusion.

We observed increased [^18^F]FDG uptake in the spinal cord (C1-T13, and thoracic T1-13) and that the uptake was higher in the cervical than the thoracic part of the mice. This suggests the importance of proximity to the lesion. In the brain, [^18^F]FDG PET has demonstrated a reduced cerebral metabolic rate for glucose after permanent MCAO and tMCAO in rats [49, 50] and mice [51]. In the spinal cord, [^18^F]FDG PET has indicated increased glycolysis and metabolic changes in EAE mice, rats with inflammatory infiltrates [37, 52, 53], transgenic Parkinson’s disease mice [54], tumor mice [55], and animal models after injuries and immune responses [56–59]. In healthy human subjects, [^18^F]FDG PET/CT showed a decreasing pattern from cervical to lumbar vertebrae and peaked at C4-6 and T12 [60–62]. In patients with acute myocardial infarction, [^18^F]FDG-PET measures of metabolic activity were significantly higher in the BM lumbar spinal cord [21, 22] and spleen and were associated with an increased risk of cardiovascular events [63, 64]. Considering the resolution (1.6-1.7 mm) of the current PET system and the size of the femur BM of the mice, we did not analyse the femur BM [^18^F]FDG uptake. In addition, due to the limited FOV of the PET scan, the lumbar part of the mice was not covered in our study.

We demonstrated reduced MSOT sO_2_ (indicating hypoxia) in the spinal cord of tMCAO mice compared to sham-operated mice. Oxygen availability varies among tissues and sites within the same tissue. Local tissue oxygen tension is one physiological regulatory mechanism of immune responses [17]. Stroke increases sympathetic nervous activity in the BM of the tMCAO mouse model [15]. tMCAO induces dynamic secondary degeneration [65, 66] and rapid activation of the peripheral immune system in the spinal cord, as well as blood–spinal cord barrier alterations in rodents [44, 46–48]. High-field magnetic resonance imaging has been used to detect oxygen metabolism in the brains of stroke patients and animal models [67–70]. Recent MSOT studies in animal models have demonstrated spinal cord hypoxia in experimental autoimmune encephalomyelitis (EAE) models [71] and white matter loss in a spinal cord injury model [72] and enabled the monitoring of image-guided stem cell therapy [73–76]. A recent study showed the feasibility of MSOT imaging of hyperoxia in the femur BM of a leukemia animal model [77]. Here, we did not observe a difference in the MSOT sO_2_ of femur BM between tMCAO mice and sham-operated mice due to the variability and small ROI size.

The spleen is a hematopoietic organ and an immediate reservoir of monocytes [9, 78]. The spleen may respond to injury in the brain by releasing stored immune cells into the blood, infiltrating into the brain and promoting a secondary inflammatory response [37]. Increased [^18^F]FDG uptake has not been reported in spleen ischemic stroke patients compared to controls [8]. Previous studies in animal models have demonstrated altered splenic function after ischemic stroke and increased levels of circulating proinflammatory cytokines [10–12, 79]. We observed no difference in splenic [^18^F]FDG uptake between tMCAO and sham-operated mice, while a positive association between [^18^F]FDG uptake in the ipsilateral striatum and spleen. In addition, we observed similar splenic sO_2_ of tMCAO mice at 24 h after occlusion compared to sham-operated mice. The splenic oxygen levels vary between 0.5 and 4.5 %, depending on the distance from the splenic artery [80].

We found a decrease in splenic mass in tMCAO mice 24 h after occlusion compared to naïve mice but not to sham-operated mice *ex vivo*. In addition, tMCAO led to a significant increase in the numbers of mature T cells in femoral BM tissues, concomitant with a stark reduction in these cell subsets in the spleen and their decrease in peripheral blood, while the numbers of mature granulocytes decreased in BM and blood and increased in the spleen. This is in line with previous reports of spleen weight reductions in mice after tMCAO [12–14], accompanied by a decreased number of monocytes/macrophages in the spleen [21, 22] and increased myeloid cells [6, 35]. Splenic atrophy was also reported in permanent MCAO in rats at 24 h [10] and at 4 days [83]. However, some studies did not include sham (surgery, but no ischemic brain injury) controls [13, 14]. Myeloid cells are trafficked to the ischemic brain, where they participate in the inflammatory response [7, 14]. Several studies have reported splenic atrophy and decreased monocytes at 4 days [12], 3-7 days [82], and 3 h-7 days [8] after tMCAO in mice. The number of mature T cells in the spleen and blood decreased and showed reduced interferon-γ production in tMCAO mice from 24 h - 14 days [84]. An earlier study showed that tMCAO increased monocyte/macrophage subsets at 3 and 7 days after ischemia in the BM, as well as an increase in circulation [14]. In addition, tMCAO activates BM HSCs and downstream hematopoietic progenitors, leading to an increased output of inflammatory monocytes and neutrophils in an animal model [15], whereas the number of lymphocyte progenitors declined. We did not observe an additional effect of cerebral ischemia on the deployment of cells from the spleen in our study. Our spleen volume and FACS data suggest that the experimental intervention dominates the egress of splenic cells into the circulation. The examination time point after occlusion and the animal model used might contribute to the different observations between studies.

In patients with acute stroke, spleen sizes were measured with abdominal ultrasound, and a statistically significant negative association was also observed between the pattern of change in total white blood cell count and spleen volume [85]. Splenic volume reduction is associated with post-stroke infection in patients [86]. Using abdominal computed tomography, another study revealed that stroke induced an initial reduction (until 48 h) followed by an increase in the splenic volume in patients [87].

There are several limitations in the current study. First, fluence correction is expected to improve the accuracy of the signal at depth [89]. Second, we only imaged the mice at 24 hours after occlusion. A longitudinal study design until several days after occlusion to monitor the dynamics will be informative about the dynamics of changes [90]. Resolution of PET and MSOT not high enough to dissect different compartments in the BM and spleen. For example, oxygen levels vary within the same tissue [80]. Here, we used the MSOT transducer array with 128 elements, and the image quality is inferior to that with 256 elements [91–93] and transmission–reflection optoacoustic ultrasound computed tomography [94]. In addition, anesthesia and mild surgical manipulation may induce granulocyte mobilization and systemic cytokine induction [88], which needs to be further elucidated.

A bidirectional relationship between hypoxia and inflammation has been demonstrated in ischemic stroke (hypoxia-induced inflammation, or inflammation reduced the rate of oxygen delivery to tissues and elicited hypoxic conditions) [95]. Further studies using molecular contrast agents (targeting hypoxia-inducible factors) [96] and transcriptomics and flow cytometry to quantify immune cells in the injured brain or spinal cord tissue will provide further mechanistic understanding.

In conclusion, we demonstrated increased metabolic activity in the spinal cord by noninvasive multimodal imaging in tMCAO mice. The inflammatory linkage among the BM, spleen, and blood after acute cerebral ischemia needs to be further investigated.

## 5. Conflict of Interest

The authors declare that the research was conducted in the absence of any commercial or financial relationships that could be construed as a potential conflict of interest.

## 6. Author Contributions

RN and JK designed the study; LM, RN, GL, SF, CA, and JK performed the experiment; RN, NS, XLDB, LM, SF, CA, and JK performed the analysis; RN and JK wrote the draft. All authors read and approved the final manuscript.

## 7. Funding

JK received funding from the Swiss National Science Foundation (320030_179277) in the framework of ERA-NET NEURON (32NE30_173678/1), the Synapsis Foundation, the Olga Mayenfisch Stiftung, and the Vontobel Foundation. RN received funding from Helmut Horten Stiftung and University of Zurich[MEDEF-20-021].

## 8. Acknowledgements

The authors acknowledge Prof. Roger Schibli, Annette Krämer at Center for Radiopharmaceutical Sciences, Department of Chemistry and Applied Biosciences, ETH Zurich; Ms Diana Kindler at Institute for Biomedical Engineering, ETH Zurich.

---

**Supplementary Table 1.**

| **Reagent** | **Source** | **Identifier** |
| --- | --- | --- |
| PBS | Gibco | 20012-019 |
| RBC Lysis Buffer 10X | BioLegend | 420301 |
| EDTA solution pH 8.0 (0.5 M) | Panreac Appli-Chem | A4892,0100 |
| Microvette 500 µL EDTA/PK100 | Sarstedt Inc | NC9990563 |
| TruStain fcX (anti-mouse CD16/32) | BioLegend | 101320 |
| CD34, FITC | Invitrogen | 11-0341-85 |
| cKIT, BV711 | BioLegend | 105835 |
| Sca1, APC-Cy7 | BD Horizon | 563991 |
| CD48, BV785 | BioLegend | 103449 |
| CD16/32, BV510 | BioLegend | 101333 |
| Lineage cocktail, PerCP-Cy5.5 | BD Pharmingen | 51-9006964 |
| CD150, PE-Cy7 | BioLegend | 115914 |
| B220, BV785 | BioLegend | 103246 |
| Gr1, PerCP-Cy5.5 | eBioscience | 45-5931-80 |
| CD4, PE-Cy7 | Invitrogen | 25-0041-82 |
| Ter119, PE | Invitrogen | 12-5921-82 |
| CD8, APC-e780 | Invitrogen | 47-0081-82 |
| CD11b, APC | eBioscience | 17-0112-82 |
| DAPI | Invitrogen | D1306 |

**Supplementary Fig. 1.**
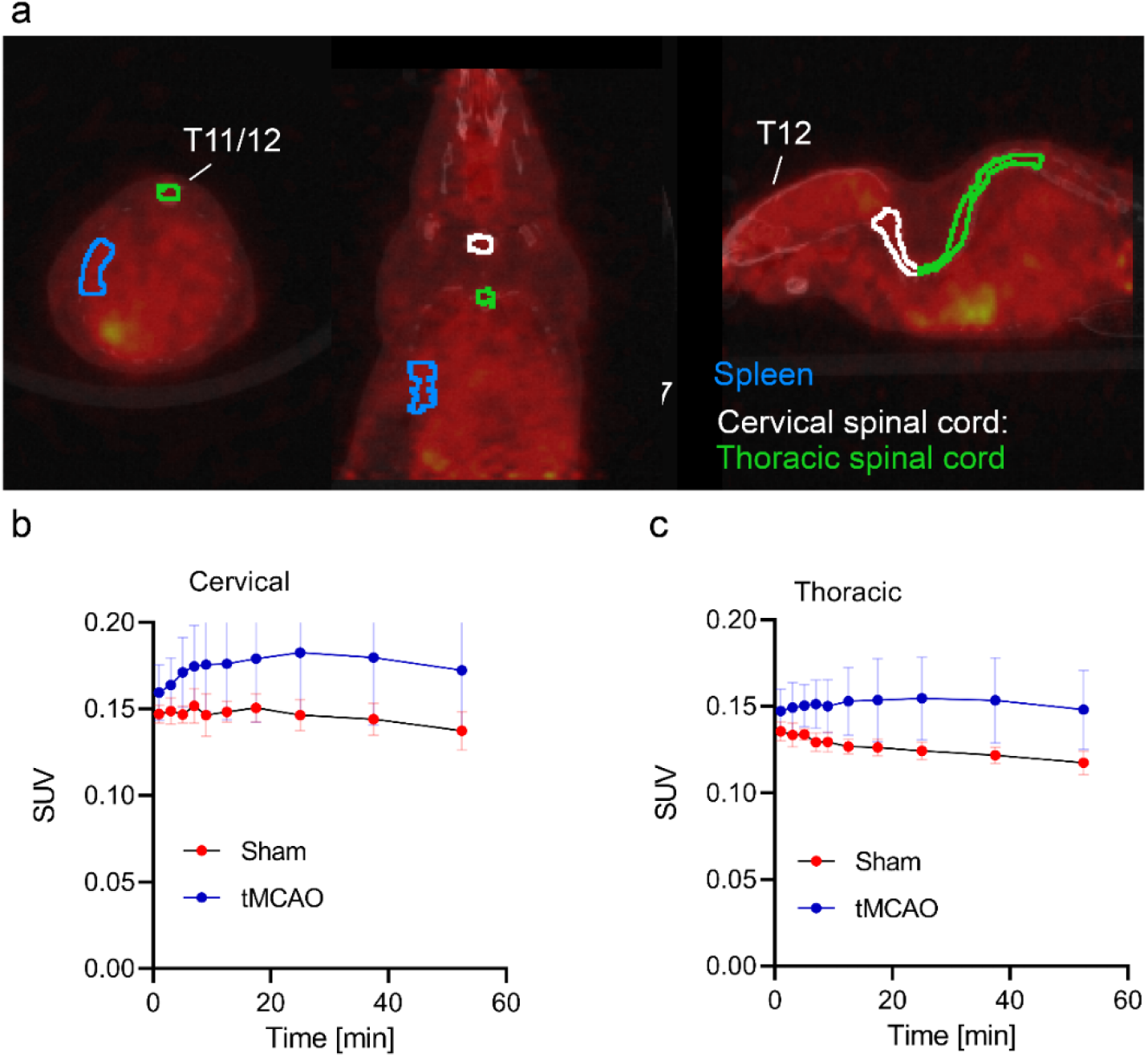
[^18^F]FDG spinal cord analysis. (**a**) Spleen (blue) spinal cord (including cervical (white) and thoracic (green)) VOI delineation on [^18^F]FDG PET. (**b**) [^18^F]FDG time activity curve in the cervical part. (**c**) [^18^F]FDG time activity curve in the thoracic part. Data are presented as the mean ± standard deviation (SD).

## Notes

### Competing Interest Statement

The authors have declared no competing interest.

